# Polo kinase phosphorylation determines *C. elegans* centrosome size and density by biasing SPD-5 toward an assembly-competent conformation

**DOI:** 10.1101/067223

**Authors:** Oliver Wueseke, David Zwicker, Anne Schwager, Yao Liang Wong, Karen Oegema, Frank Jülicher, Anthony A. Hyman, Jeffrey B. Woodruff

## Abstract

Centrosomes are major microtubule-organizing centers composed of centrioles surrounded by an extensive proteinacious layer called the pericentriolar material (PCM). In *C. elegans* embryos, the mitotic PCM expands by Polo-kinase (PLK-1) phosphorylation-accelerated assembly of SPD-5 molecules into supramolecular scaffolds. However, how PLK-1 phosphorylation regulates SPD-5 assembly is not known. We found that a mutant version of SPD-5 that is insensitive to PLK-1 phosphorylation (SPD-5^4A^) could localize to PCM but was unable to rescue the reduction in PCM size and density when wild-type SPD-5 levels were decreased. In vitro, purified SPD-5^4A^ self-assembled into functional supramolecular scaffolds over long time scales, suggesting that phosphorylation only controls the rate of SPD-5 scaffold assembly. Furthermore, the SPD-5 scaffold, once assembled, remained intact and supported microtubule nucleation in the absence of PLK-1 activity *in vivo*. We conclude that Polo Kinase is required for rapid assembly of the PCM scaffold but not for scaffold maintenance or function. Based on this idea, we developed a theoretical model that adequately predicted PCM growth rates in different mutant conditions *in vivo*. We propose that PLK-1 phosphorylation-dependent conversion of SPD-5 into an assembly-competent form underlies PCM formation *in vivo* and that the rate of this conversion determines final PCM size and density.

## INTRODUCTION

Centrosomes are the main microtubule-organizing centers of animal cells, consisting of a pair of centrioles that organize a dynamic protein mass called pericentriolar material (PCM). The PCM consists of a structured, small interphase layer [1,2] around which, amorphous mitotic PCM assembles to achieve maximal microtubule nucleation to facilitate mitotic spindle assembly. Despite the importance of PCM in microtubule organization, the mechanisms behind mitotic PCM formation remain elusive. In *C.elegans*, large-scale RNAi screens and genetics have revealed a limited number of components required for PCM assembly: SPD-2 [3,4], the polo-like kinase PLK-1 [5] and SPD-5 [6]. It was also shown that similar proteins are involved in PCM assembly in other species, suggesting that a universal assembly mechanism may exist. For instance, PCM assembly in vertebrates and *Drosophila* requires the assembly of large proteins such as Pericentrin/D-PLP and Cdk5RAP2/Centrosomin, which resemble SPD-5 in that they contain numerous interspersed coiled-coil domains [6–11]. The assembly of these proteins is facilitated by SPD-2/Cep192 and the phosphorylation activity of the conserved Polo-like-kinase Plk1/PLK-1 [3–5, 12–15]. However, how these molecular interactions lead to PCM assembly and determine final PCM size and density remain outstanding questions.

We previously hypothesized that PCM is nucleated at centrioles and then rapidly expands via autocatalytic incorporation of cytosolic PCM components [16]. In our model, unassembled PCM proteins exist in a soluble form that can transition into an assembly-competent state within the PCM and then become stably incorporated. Once incorporated, PCM proteins will recruit additional PCM components, which is an autocatalytic event. Consequentially, in our model the kinetics of PCM assembly depend on the rate by which PCM material, after being recruited to the centrosome, converts from the soluble to the assembly-competent form. The existence of such a conversion is supported by the observations that *C. elegans* PCM proteins are indeed monomeric in cytoplasm prior to assembly, whereas they interact at the centrosome [17].

We recently reported that purified SPD-5 can form supramolecular PCM-like assemblies *in vitro*, the formation of which is accelerated by PLK-1 phosphorylation of SPD-5 [12]. Mass spectrometry revealed PLK-1 phosphorylation sites on SPD-5 that, when mutated, prevented PCM assembly *in vivo*. These results indicate that PCM formation in *C. elegans* is driven by Polo Kinase-mediated oligomerization of SPD-5 around centrioles. We also found that only SPD-5 assemblies, and not SPD-5 monomers, recruited other PCM proteins, including PLK-1, leading us to propose that this emergent scaffolding property of SPD-5 could be the basis for autocatalytic PCM expansion in vivo [12,17]. Additionally, our vitro experiments revealed that the stability and scaffolding capacity of SPD-5 assemblies were independent of PLK-1 phosphorylation, suggesting that Polo Kinase activity may only regulate the speed of SPD-5 assembly. However, we did not test whether Polo Kinase has additional roles in PCM maintenance or function *in vivo*. Nor did we test if unphosphorylated SPD-5 can be recruited to existing PCM and then be converted to an assembly-competent state as predicted by our previous model [16].

In this study, we combined *in vivo* analysis, *in vitro* reconstitution, and modeling to investigate how Polo Kinase regulates SPD-5 assembly to form PCM in *C. elegans*. Our results indicate that Polo Kinase phosphorylation affects the rate of SPD-5 assembly without dramatically affecting SPD-5 recruitment to existing PCM, PCM stability, or PCM function. Additionally, we conclude that a phospho-site binding mechanism cannot explain PCM assembly. Rather, we propose that SPD-5 naturally isomerizes between assembly-incompetent and assembly-competent states, and that Polo Kinase phosphorylation biases SPD-5 toward the latter state. Furthermore, our results suggest that the conversion rate of SPD-5 into an assembly competent state is the key determinant of final PCM size and density *in vivo*.

## RESULTS AND DISCUSSION

### A SPD-5 phospho-mutant binds to PCM in *C. elegans* embryos

We first set out to determine if PLK-1 phosphorylation of SPD-5 is required only for PCM expansion or also for SPD-5 binding to PCM. For this purpose we used spinning disk confocal microscopy to observe PCM assembly in *C. elegans* embryos expressing GFP-tagged versions of SPD-5 (SPD-5^WT^ and SPD-5^4A^) as their sole source of SPD-5. SPD-5^4A^ is mutated in the critical sites for PLK-1 mediated centrosome assembly *in vivo* (Figure 1A; [12]). As previously shown, GFP::SPD-5^4A^ localized to pre-mitotic centrosomes, but, in contrast to GFP::SPD-5^WT^, failed to expand the PCM (Figure 1B; [12]). To test if SPD-5^4A^ is still capable of binding to existing PCM, we observed PCM assembly in embryos expressing endogenous SPD-5 and GFP::SPD-5^4A^. In such embryos, mitotic PCM assembled and GFP::SPD-5^4A^ localized to the PCM (Figure 1B).

**Fig. 1.**
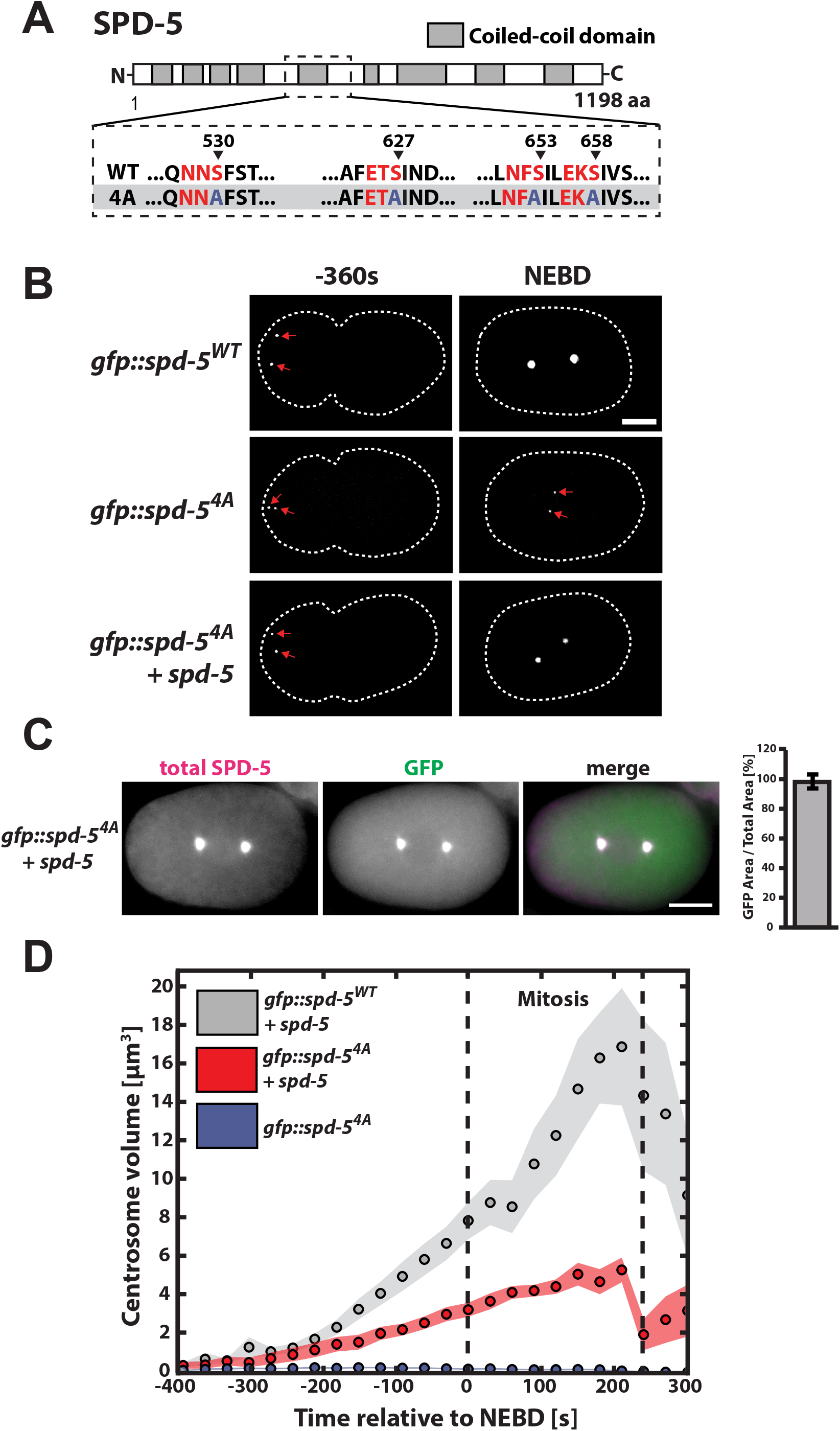
A SPD-5 phospho-mutant binds to PCM and reduces PCM size in *C. elegans* embryos. **(A)** Diagram of SPD-5^WT^ and SPD-5^4A^ sequence. Canonical PLK-1 consensus motifs [25,26] are indicated in red. The arrowheads indicate the phosphorylated residues in each motif. **(B)** Embryos expressing GFP::SPD-5^WT^ or GFP::SPD-5^4A^ in the absence or presence of endogenous SPD-5 were imaged during the progression of the first cell cycle. Images show maximum projections of embryos prior to and at nuclear envelope breakdown (NEBD). Dashed white line indicates cell outline. Scale bar, 10 μm. **(C)** Embryos expressing endogenous SPD-5 and GFP::SPD-5^4A^ were fixed and stained for SPD-5 and GFP. Centrosome areas were determined from total SPD-5 and GFP::SPD-5^4A^ signal for embryos prior to or at mitotic onset. Overlap between total SPD-5 and GFP::SPD-5^4A^ was estimated to be 98 ± 5% (mean ± SEM, n = 5). Scale bar, 10 μm. **(D)** Centrosome volumes were determined from centrosome area measurements for embryos expressing GFP::SPD-5^WT^ (n = 8) or GFP::SPD-5^4A^ (n = 9) in the presence of endogenous SPD-5, as well as embryos expressing only GFP::SPD-5^4A^ (n = 10). Circles represent mean values and shaded areas represent SEM.

### PCM size and density are reduced in embryos ectopically expressing a GFP::SPD-5^4A^ transgene

Quantification and comparison of centrosome area from immunostainings against SPD-5 and GFP revealed perfect overlap of total SPD-5 and GFP::SPD-5^4A^ signals, showing that GFP::SPD-5^4A^ localizes throughout the entire PCM and can be used to determine PCM size in embryos expressing mutant SPD-5 (Figure 1C). By comparing centrosome sizes determined from GFP signal we found that PCM assembled in the presence of endogenous SPD-5 and GFP::SPD-5^4A^ was ~58% smaller in volume (3.8 ± 0.4 μm^3^, mean ± SEM) at nuclear envelope breakdown (NEBD) than PCM assembled with endogenous SPD-5 and wild-type GFP::SPD-5 (8.0 ± 1.0 μm^3^, mean ± SEM) (Figure 1D). As reported previously, GFP::SPD-5^WT^ and GFP::SPD-5^4A^, as well as endogenous SPD-5 levels, are similar in these worms (~30% transgene, ~70% endogenous), suggesting that this difference does not result from altered SPD-5 concentrations (Figure S1A, [12]). We conclude that SPD-5^4A^ binds to PCM and that its expression reduces PCM expansion.

In addition to being smaller, centrosomes assembled in the presence of the SPD-5^4A^ mutant were also less dense. Based on the assumption that SPD-5 forms the underlying PCM scaffold [6,12], we used GFP::SPD-5 fluorescence at the PCM to approximate PCM density from the mean pixel intensity of maximum intensity z-projections. In embryos expressing endogenous SPD-5 and GFP::SPD-5^WT^, centrosomal GFP signal increased with time after fertilization until the onset of mitosis (Figure 2A, gray points), indicating an increase in PCM density up to mitosis. In embryos expressing endogenous SPD-5 and GFP::SPD-5^4A^, centrosomal GFP signal started at a similar mean intensity shortly after fertilization but remained constant until mitosis (Figure 2A, red points). A comparison of GFP fluorescence of centrosomes at NEBD revealed 26% higher intensity for centrosomes assembled with GFP::SPD-5^WT^ (WT = 420 ± 99 a.u. vs. 4A = 332 ± 76 a.u., mean ± STD; Figure 2B). We tested if this difference in GFP intensity resulted from hampered binding of GFP::SPD-5^4A^ to PCM or generally a lower SPD-5 density at the PCM. Immunostainings against GFP and SPD-5 showed that SPD-5 levels at the PCM were reduced in embryos expressing the 4A mutant (Figure 2C and D) and that the ratios of transgenic GFP::SPD-5 to total SPD-5 immunostaining signal at wild-type and mutant PCM were very similar (Figure S2A). These results suggest that the reduction in GFP fluorescence seen in GFP::SPD-5^4A^ embryos reflects a general reduction of SPD-5 levels at the PCM. Also, immunostainings revealed a similar difference in SPD-2 and PLK-1 levels at wild type and mutant PCM (Figure 2E and 2F), indicating that concentrations of SPD-2 and PLK-1 correlate with SPD-5 concentration at the PCM. Thus, expression of GFP::SPD-5^4A^ reduces volume and density of the functional PCM scaffold.

**Fig. 2.**
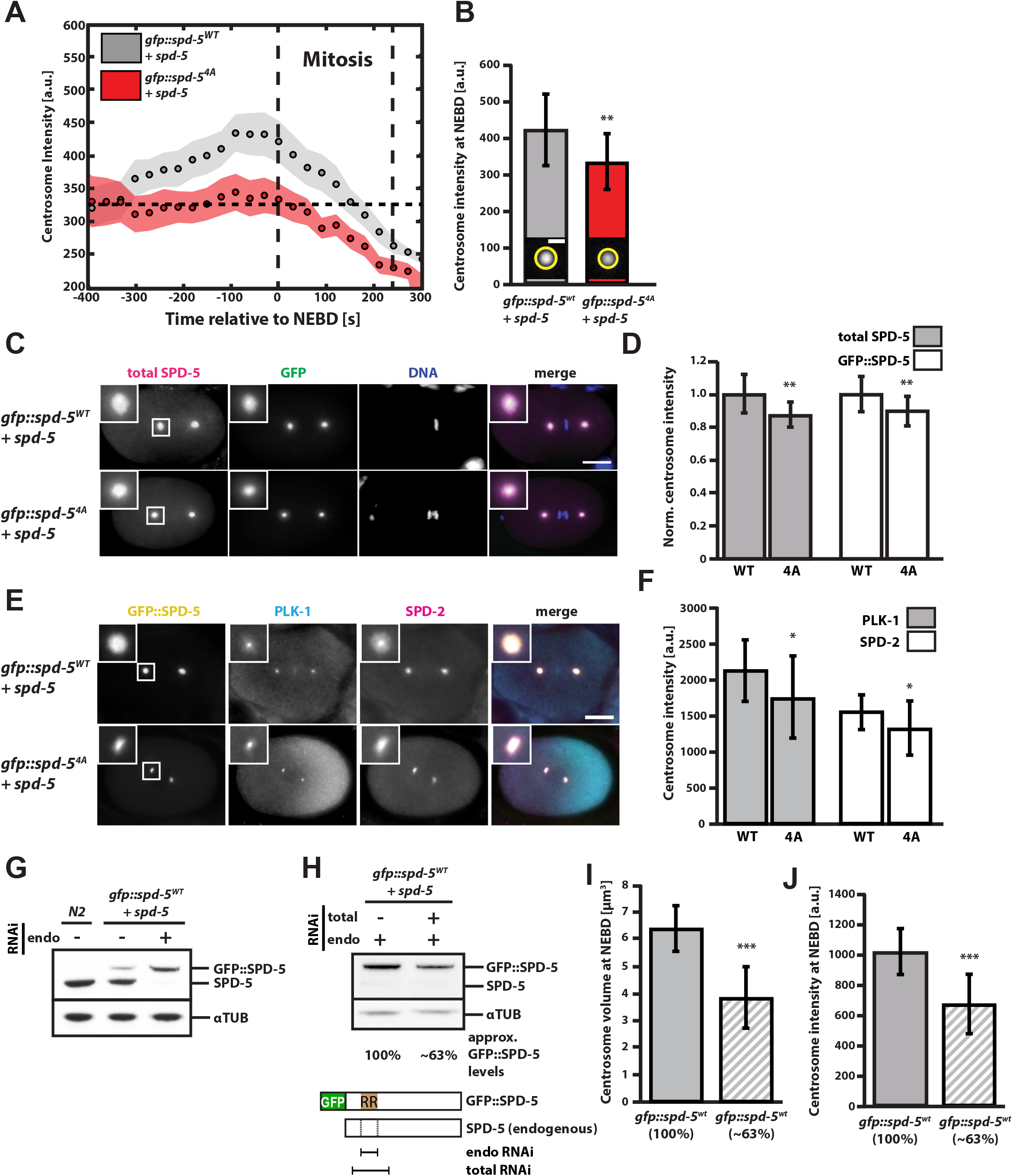
The concentration of phosphorylation-receptive SPD-5 determines PCM size and density *in vivo*. **(A)** Mean maximum GFP intensity of centrosomes in embryos expressing endogenous SPD-5 and GFP::SPD-5^WT^ (n = 10) or endogenous SPD-5 and GFP::SPD-5^4A^ (n = 10) over time relative to NEBD. Circles represent mean values and shaded areas represent SEM. **(B)** Mean maximum GFP intensity of centrosomes at NEBD as in (A) showing a 26% difference in intensity of centrosomes assembled in the presence of GFP::SPD-5^WT^ or GFP::SPD-5^4A^. Bars indicate mean ± STD (**, p<0.01). **(C)** Embryos expressing GFP::SPD-5^WT^ plus endogenous SPD-5 (n = 17) or GFP::SPD-5^4A^ plus endogenous SPD-5 (n = 19) were fixed and immunostained for SPD-5 and GFP. Scale bar represents 10 μm. **(D)** Ratio of GFP and SPD-5 immunofluorescence signal from (C). Wild type signals were used for normalization. Bars represent mean ± STD (**, p<0.01). **(E)** Embryos expressing endogenous SPD-5 and GFP::SPD-5^WT^ or GFP::SPD-5^4A^ were fixed and stained for SPD-2 and PLK-1. Insets show close ups of centrosomes. Scale bar, 10 μm. **(F)** Quantification of **(E)** using centrosomal SPD-2 and PLK-1 immunofluorescence signal in GFP::SPD-5^WT^ (n = 19) or GFP::SPD-5^4A^ (n = 25) containing centrosomes. Bars represent mean ± STD (*, p<0.05). **(G)** Western blot showing the compensation of SPD-5 levels upon expression of the transgenic or knock down of endogenous SPD-5. RNAi was carried out for 24 hr specifically against endogenous SPD-5. **(H)** Western blot showing full depletion of endogenous SPD-5 and partial depletion of transgenic SPD-5 after 12 hrs of mixed RNAi treatment. Quantification of the SPD-5 levels showed that mixed RNAi treatment lowered the transgenic SPD-5 levels to ~63% of the control situation where only endogenous SPD-5 was depleted. The schematic shows the target sequences of RNAi constructs targeting endogenous SPD-5 (endo RNAi) or endogenous and transgenic SPD-5 (total RNAi). **(I)** Centrosome volumes at NEBD in embryos expressing GFP::SPD-5^WT^ exclusively (100%, n = 19) or reduced levels of GFP::SPD-5^WT^ (~63%, n = 23). Bars represent mean ± STD (***, p<0.001). **(J)** As in **(H)** but showing mean maximum centrosomal GFP fluorescence at NEBD. Bars represent mean ± STD (***, p<0.001).

### The concentration of phosphorylation-receptive SPD-5 determines PCM size and density *in vivo*

How can the presence of a mutated transgenic SPD-5 cause a reduction of PCM volume and density? It is possible that GFP::SPD-5^4A^ acts as a dominant negative mutant that interferes with accumulation of wild-type SPD-5 at the PCM, possibly by occupying and blocking required SPD-5 binding sites in the PCM scaffold. Alternatively, GFP::SPD-5^4A^ may act as a loss-of-function mutant, and, due to protein level compensation, the phenotype seen in embryos expressing mutant SPD-5 could be a consequence of the reduction of available wild-type SPD-5. We previously observed such compensation of the centrosomal protein SPD-2; however, ectopic expression of a codon-adapted version of SPD-5::GFP did not influence the expression of endogenous SPD-5 [5]. Surprisingly, ectopic expression of transgenic GFP::SPD-5^WT^ in our current strain led to a reduction in endogenous SPD-5, and selective RNAi against endogenous SPD-5 lead to an upregulation of transgenic SPD-5 (Figure 2G). These different behaviors could be caused by the sequence differences in the SPD-5 transgenes. The GFP::SPD-5^WT^ transgene used in this study is codon adapted only at the N-terminus (Woodruff 2015), while the SPD-5::GFP transgene used in Decker et al. (2011) was codon adapted throughout its sequence. Thus, codon optimization of SPD-5 interferes with regulation of total SPD-5 levels. We conclude that SPD-5 levels are normally tightly regulated and that worms expressing the transgenic SPD-5 used in this study compensate by down-regulating endogenous wild-type SPD-5.

To test if PCM volume and density respond to SPD-5 concentration changes, we fully removed endogenous SPD-5 and then reduced the concentration of GFP::SPD-5^WT^ using a double RNAi condition targeting the endogenous and transgenic *spd-5* transcripts with different strengths. Using this method, we reduced GFP::SPD-5^WT^ levels to about 63% compared to the control condition, while fully removing the endogenous copy in both cases (Figure 2H). Reduction of GFP::SPD-5^WT^ reduced PCM volume by ~40% (3.9 ± 1.2 μm^3^, p < 0.001) and PCM density by ~34%, (p < 0.001) (Figure 2I and 2J). These changes in centrosome size and density were similar to changes observed when GFP::SPD-5^4A^ was ectopically expressed (see Figure 1D and 2A). These results are consistent with GFP::SPD-5^4A^ being a loss-of-function mutant. Furthermore, we conclude that the concentration of available wild-type SPD-5 determines PCM size and density.

### PLK-1 phosphorylation of SPD-5 affects the rate of PCM matrix assembly without affecting matrix function *in vitro*

Our *in vivo* analysis showed that PCM size and density are reduced in embryos when either the concentration of wild-type SPD-5 is reduced via RNAi or when SPD-5^4A^ is ectopically expressed in addition to endogenous SPD-5. However, these experiments did not allow us to exclude the possibility that SPD-5^4A^ could act as a dominant negative mutant. We directly tested this hypothesis using an *in vitro* assay for PCM assembly developed in our lab [12].

Similar to our observations *in vivo*, we found that purified SPD-5^4A^::GFP localized to PCM-like networks formed with SPD-5^WT^::TagRFP *in vitro* (Figure 3A). Next, we tested the effect of SPD-5^4A^::GFP on wild-type SPD-5 network growth in the presence of PLK-1. We prepared network reactions on ice with equimolar amounts of SPD-5^WT^::GFP and PLK-1, then added either buffer (WT), SPD-5^4A^, (WT+4A), or additional SPD-5^WT^ (WT+WT). We warmed the tubes to 23°C to initiate network assembly, then, after 30 min, we squashed a sample under a cover slip for analysis. Under these conditions, small, nascent networks could be seen in the control sample (Figure 3B), and we verified that growth had not yet plateaued (unpublished data); thus, our experiments should allow detection of any stimulatory or inhibitory effects.

**Fig. 3.**
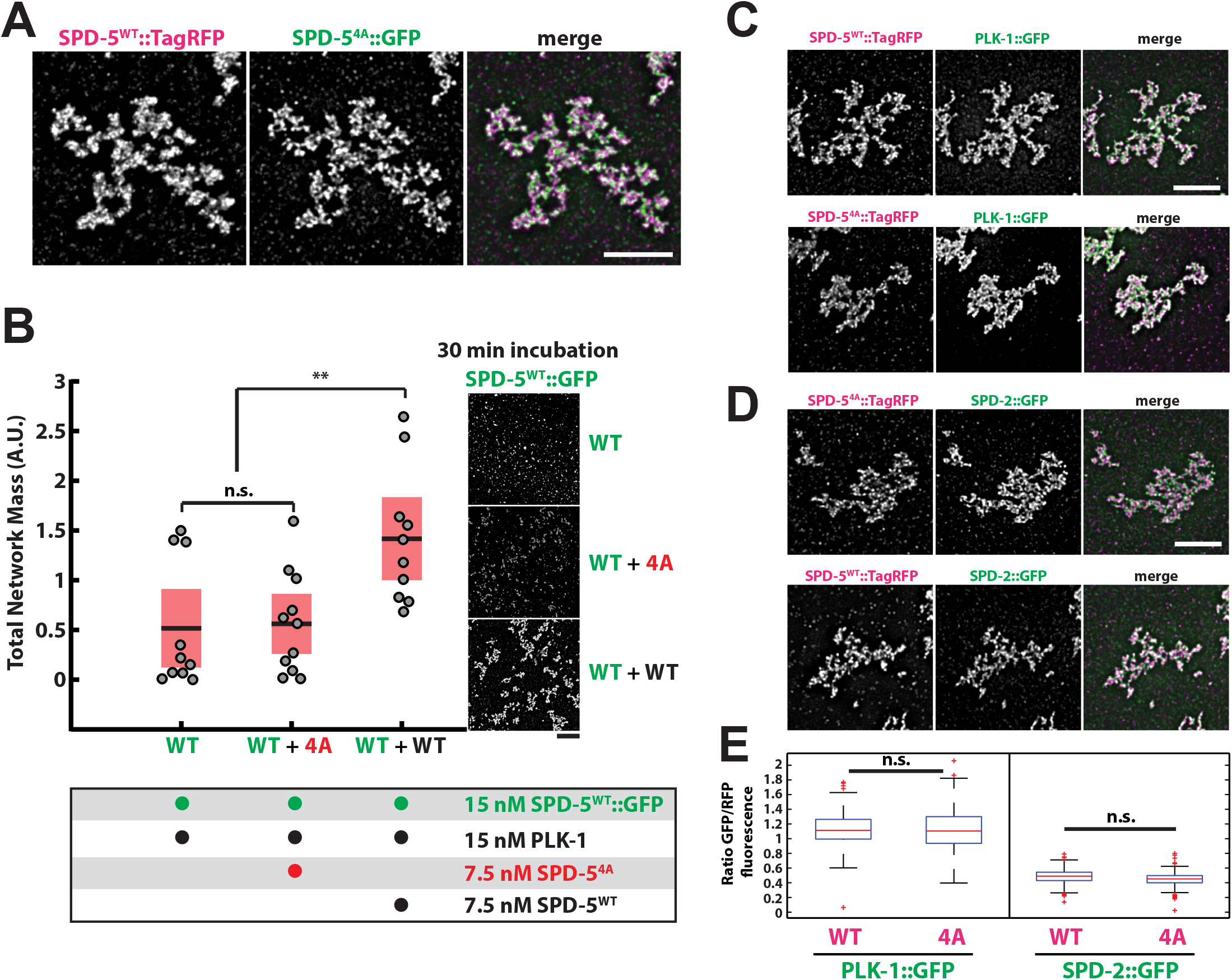
PLK-1 phosphorylation of SPD-5 affects the rate of PCM matrix assembly without affecting matrix function *in vitro*. **(A)** 25 nM SPD-5^WT^::TagRFP was mixed with 25 nM SPD-5^4A^::GFP to assemble SPD-5 networks as previously described [10]. Networks were squashed under a coverslip and imaged using fluorescence microscopy. Scale bar, 5 μm. **(B)** Quantification of total SPD-5^WT^::GFP network mass after 30 min. Networks were assembled from 15 nM SPD-5^WT^::GFP and 15 nM PLK-1 supplemented with buffer (WT, n=10), 7.5 nM SPD-5^4A^::TagRFP (WT+4A, n=11), or 7.5 nM SPD-5^WT^::TagRFP (n = 10). Horizontal black bars represent the mean, and red shaded areas represent 95% confidence intervals (n.s., p =0.87; **, p<0.01). Scale bar, 5 μm. **(C)** SPD-5 networks were assembled from 25 nM SPD-5^WT^::TagRFP or 25 nM SPD-5^4A^::TagRFP and incubated with 25 nM PLK-1::GFP. Scale bar, 5 μm. **(D)** Same as **(C)** but incubated with 25 nM SPD-2::GFP. Scale bar, 5 μm. **(E)** Quantification of PLK-1::GFP and SPD-2::GFP fluorescence from (C) and (D). Mean values are indicated with red lines (n = 148 networks per condition; n.s., p>0.10).

Total network mass was ~2-fold higher in the sample containing unlabeled SPD-5^WT^ compared to the control where only buffer was added (WT+WT vs. WT; p = 0.007) (Figure 3B). In contrast, total network mass in the sample containing SPD-5^4A^ was only slightly higher than the control sample, and the difference was not statistically significant (p = 0.86) (Figure 3B). These data indicate that during PCM assembly SPD-5^4A^ behaves as a loss-of-function mutant rather than a dominant-negative mutant. Furthermore, our *in vitro* results corroborate our *in vivo* findings that PCM assembly rate is largely determined by the amount of phosphorylation-responsive SPD-5 available in the system.

We then used this *in vitro* assay to test if PLK-1 is also required for proper functioning of the PCM scaffold. As observed previously, SPD-5^4A^ assembled into supramolecular networks at a rate similar to unphosphorylated wild-type protein [12]. After one hour of incubation at 23°C, networks exclusively assembled from SPD-5^WT^::TagRFP or SPD-5^4A^::TagRFP equivalently recruited SPD-2::GFP and PLK-1::GFP (Figure 3C-E), suggesting that SPD-5^4A^ and unphosphorylated SPD-5^WT^ can form functional PCM scaffolds *in vitro* given sufficient time.

### PLK-1 phosphorylation is not required to maintain PCM scaffold stability or function *in vivo*

Our *in vitro* results predict that SPD-5 scaffolds, once formed, should function without needing continuous PLK-1 phosphorylation *in vivo*. To test this idea, we constructed a *C. elegans* strain expressing GFP::SPD-5^WT^ and an analog-sensitive PLK-1 mutant (PLK-1^AS^) that can be inhibited by the drug 1NM-PP1 *(plk1Δ; plk-1^as^ gfp::spd-5;* [21]). We permeabilized embryos using partial knockdown of *perm-1* via RNAi [22], then identified pre-mitotic embryos where centrosomes had formed but were not yet full-sized. Addition of 10 μM 1NM-PP1 to these embryos arrested centrosome growth: both centrosome size and GFP::SPD-5 fluorescence remained constant thereafter (Figure 4A and 4B; n = 10). However, centrosomes continued to grow if DMSO was added instead (Figure 4C; n = 10). Thus, PLK-1 is not required to maintain SPD-5 at the centrosome; this stands in stark contrast to gamma tubulin, which does require continuous PLK-1 activity for centrosomal localization [12].

**Fig. 4.**
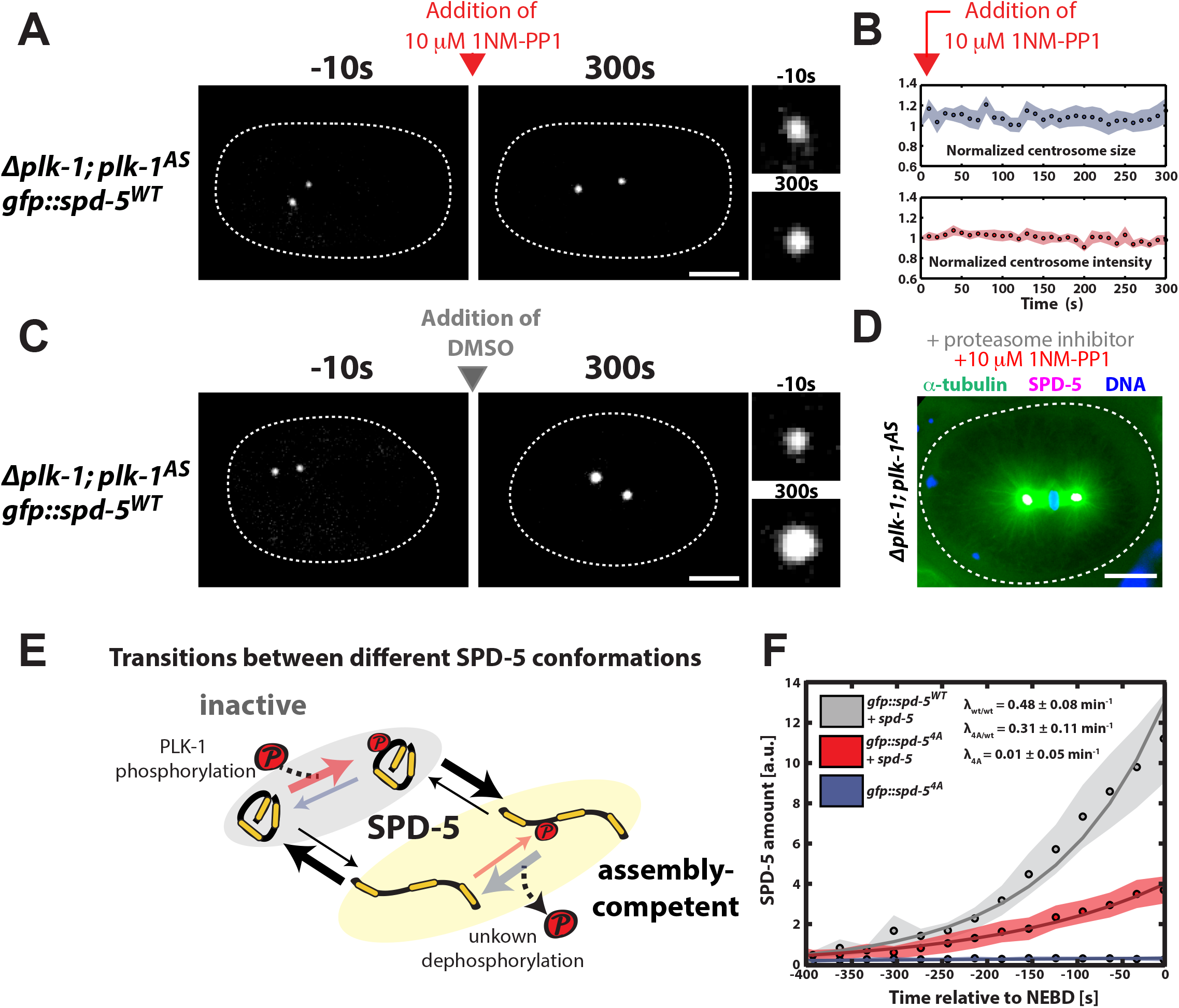
A PLK-1 dependent SPD-5 conversion model can explain *in vivo* PCM assembly. **(A)** *plk-1^AS^* embryos expressing GFP::SPD-5^WT^ were visualized by fluorescence confocal microscopy (white dashed line is embryo outline). 10 μM 1-NM-PP1 (PLK-1^AS^ inhibitor) was added to permeabilized embryos prior to mitotic entry (red arrow). Insets on the right represent zoomed in images of a centrosome. **(B)** Quantification of centrosome size and fluorescence intensity of GFP::SPD-5^WT^ at centrosomes from (A) (n = 10 centrosomes). Circles represent mean values and shaded areas represent SEM. **(C)** Same as in (A), except that DMSO was added instead of the PLK-1^AS^ inhibitor. Centrosomes continued to grow as expected. **(D)** *plk-1^AS^* embryos were treated with 10 μM 1-NM-PP1 and 20 μM c-lactocystin-β-lactone for 20 min to arrest cells in metaphase, then frozen and fixed in methanol. Tubulin, SPD-5 and DNA were visualized by immunofluorescence. Centrosomes continued to nucleate microtubules in the absence of PLK-1 activity. **(E)** Possible transitions between the different conformations of SPD-5. **(F)** Total SPD-5 amount fit with the accumulation model to determine SPD-5 incorporation rates. Circles represent mean values and shaded areas represent SEM from experimental data. Solid lines represent the fits.

To test the functionality of PCM-localized SPD-5 in the absence of PLK-1 phosphorylation *in vivo*, we treated permeabilized embryos with 10 μM 1NM-PP1 and the proteasome inhibitor c-lactocystin-β-lactone for 20 min, then fixed the embryos and visualized microtubules using immuofluorescence. Centrosomes still nucleated microtubules and formed spindles after PLK-1 inhibition, suggesting that SPD-5 retains its functional capacity for scaffolding in the absence of PLK-1 phosphorylation (Figure 4D; n = 8). These *in vivo* results are in agreement with our *in vitro* data and suggest that PLK-1 phosphorylation is not required for the maintenance or function of SPD-5 scaffolds but instead only controls the rate of SPD-5 scaffold formation, and, subsequently, PCM assembly.

### A PLK-1-dependent SPD-5 conversion model can explain *in vivo* PCM assembly

Our data, combined with previous studies, allow us to propose a simple mechanism for PLK-1-dependent PCM assembly in *C.elegans*. Prior to incorporation into the PCM, SPD-5 is mostly monomeric and does not interact with SPD-2 or PLK-1 [17]. We term this the inactive form of SPD-5, which cannot contribute to PCM assembly itself but can localize to centrioles and segregate into existing PCM (Figure 4E). Since GFP::SPD-5^4A^ alone is not capable of expanding PCM *in vivo* (Figure 1B and D), we assume that SPD-5^4A^ as well as unphosphorylated SPD-5^WT^ exist primarily in the inactive form. Secondly, we define an active, assembly-competent form, which can self-assemble into supramolecular structures and contribute to PCM growth (Figure 4E). Because our *in vitro* data show that purified SPD-5 can spontaneously self-assemble and that PLK-1 phosphorylation accelerates this assembly process [12], we propose that SPD-5 can transition into the assembly-competent state spontaneously and that this transition is much more likely if SPD-5 is phosphorylated by PLK-1. Thus, we assume that Polo kinase-phosphorylated SPD-5 exists predominately in the assembly-competent state.

Based on this idea we constructed a mathematical model (see SI) in which the inactive form of SPD-5 can be converted locally at the centrosome into the assembly-competent form through an active process such as Polo kinase phosphorylation [16]. We used this model to fit the accumulation rate of total SPD-5 at centrosomes with an exponential function to describe the rate of PCM growth (see SI). Total amounts of SPD-5 were estimated from centrosome volumes multiplied by SPD-5 densities (Figure S3A). We fit the accumulation rate of total SPD-5 from initiation of assembly until NEBD (Figure 4F). When embryos only expressed SPD-5^WT^, the PCM growth rate was 0.48 ± 0.08 min^−1^. In contrast, when embryos only expressed SPD-5^4A^, PCM growth rate was only 0.01 ± 0.05 min^−1^. We then used these measured rates to predict the PCM growth rate in the mixed scenario where SPD-5^4A^ is present in a background of endogenous SPD-5^WT^. Based on western blot analysis, we estimated that ~70% of SPD-5 protein is wild-type and ~30% is mutated in these embryos (Figure S1A). Using these values, our model predicted that the PCM accumulation rate in these embryos should be 0.34 ± 0.06 min^−1^ (see SI for calculation details). This value is very similar to the accumulation rate we obtained when fitting the data (0.31 ± 0.11 min^−1^). Taken together, these results suggest that a model based on centrosomal conversion of SPD-5 into an assembly-competent form is adequate to describe the complex process of PCM assembly in *C.elegans* embryos.

How does PLK-1 change SPD-5 to induce self-assembly? *In vitro* both unphosphorylated SPD-5^WT^ and SPD-5^4A^ assembled into supramolecular networks after a long period of time (Figure 3 and [12]); but, addition of PLK-1 dramatically accelerated SPD-5^WT^ self-assembly. These results suggest that SPD-5 naturally isomerizes between the inactive form and the assembly-competent form, and that PLK-1 phosphorylation of SPD-5 lowers the energy barrier of this transition (Figure S3B). The role of PLK-1 in PCM assembly, then, is to bias SPD-5 isomerization towards the assembly-competent state. Besides PLK-1 phosphorylation, PCM assembly requires SPD-2, a protein known to control centrosome growth rate and thus size *in vivo* [5,23]. We have shown previously that SPD-2 accelerates SPD-5 assembly in the presence and absence of PLK-1 *in vitro*, demonstrating that multiple mechanisms regulate SPD-5 self-assembly [12]. Whether SPD-2 also enhances SPD-5 self-assembly by affecting its isomerization or by some other process remains to be investigated. We conclude that SPD-5 has the intrinsic capability to assemble functional PCM and that PLK-1 phosphorylation and SPD-2 simply accelerate the rate of assembly.

An unexpected observation from our experiments was that PCM density increased over time from fertilization until mitosis in wild-type embryos. However, when we reduced the speed of PCM assembly by reducing the available pool of phosphorylation-receptive SPD-5, PCM density did not change over time. We do not understand the molecular basis of this phenomenon. One possibility is that this is caused by a buildup of elastic stress as the centrosome grows. If the growth rate is faster than the stress relaxation rate, centrosome density will increase. However, if the growth rate is slowed down, for instance, by reducing the concentration of wild-type SPD-5, then the stress could relax and centrosome density would remain constant. This supports the idea that the PCM is not solid, but rather a viscous, gel-like material, which forms by phase transition of soluble molecules into PCM. This notion is further supported by the observation that GFP::SPD-5^4A^ localized to PCM without interfering with the expansion of the scaffold.

In *D. Melanogaster*, PCM assembly is driven by Polo kinase-regulated multimerization of the scaffolding protein Centrosomin [13], suggesting that Centrosomin is the functional homolog of SPD-5. Similar to SPD-5, Centrosomin must be phosphorylated at multiple residues to achieve its full scaffolding potential [13]. In vertebrate cells, Polo kinase and the Centrosomin homolog CDK5Rap2 are also required for mitotic PCM assembly [24,25]. Thus, a common mechanism for PCM assembly is emerging that centers on Polo kinase-mediated phosphorylation of large coiled-coil proteins. It will be of interest to determine the similarities in self-assembly properties and regulation of SPD-5, Centrosomin, and CDK5Rap2.

## MATERIALS AND METHODS

### Worm strains

*C. elegans* worm strains were maintained following standard protocols [26]. For this study we used previously described worm strains OD847 *(gfp::spd-5^WT^)* and OD903 *(gfp::spd-5^4A^)* [12]. Briefly, both strains contain MosSCI single-copy integrants of *gfp::spd-5* transgenes on Chromosome II rendered RNAi-resistant by re-encoding the sequence between nucleotides 500 to 1079 in the *spd-5* genomic sequence. Their genotypes are as follows:

OD847: unc-119(ed9) III; ltSi202[pVV103/ pOD1021; Pspd-2::GFP::SPD-5 RNAi-resistant; cb-unc-119(+)]II

OD903: unc-119(ed9) III; ltSi228[pVV153/ pOD1615; Pspd-2::GFP::spd-5 S530A, S627A, S653A, S658A reencoded; cb-unc-119(+)]II

We also used two new lines expressing *gfp::spd-5* with *plk-1^WT^* (OD2420) / *plk-1^AS^* (OD2421) in a *plk-1* deletion background. The genomic *plk-1* locus *(plk-1^WT^)* was amplified and the analog sensitive Shokat allele, *plk-1^AS^*, with the C52V and L115G mutations was generated using site-directed mutagenesis. *plk-1^WT^* and *plk-1^AS^* transgenes were crossed into a *plk-1* deletion background. A *gfp::spd-5^WT^* transgene was subsequently crossed into each resultant strain to allow direct monitoring of PCM. Their genotypes are as follows:

OD2420: unc-110(ed9) plk-1(ok1787) III; ltSi654[pOD1021; pspd-2::GFP::spd-5 RNAi-resistant; cb-unc-119(+)I; ltSi54[pOD1042; Pplk-1::PLK-1; cb-unc-119(+)II

OD2421: unc-110(ed9) plk-1(ok1787) III; ltSi654[pOD1021; pspd-2::GFP::spd-5 RNAi-resistant; cb-unc-119(+)I; ltSi55[pOD1048; Pplk-1::PLK-1 C52V, L115G; cb-unc-119(+)II

#### RNAi treatments

RNAi against endogenous *spd-5* was carried out as previously described [27]. Briefly, the following primers were used to amplify nucleotides 501 – 975 of endogenous *spd-5* from cDNA and cloned into Gateway^®^ pDonor^™^221 vector via BP reaction to create spd-5-pENTR^TM^ vector: *spd-5-fw* (GGG GACAAGTTT GTACAAAAAAGCAG GCT ggaattgtccgctactgatg), *spd-5-rev* (GGGACCACTTTGTACAAGAAAGCTGGGTgtgctcaagcttgctacac). The amplified sequence was then transferred to L4440_GW (Addgene) destination vector and used to transform HT115(DE3) bacteria strain for RNA expression. Using this feeding clone, full knock down of endogenous SPD-5 was achieved typically within 24 hs. Simultaneous RNAi against endogenous and transgenic SPD-5 was carried out using the SPD-5 (F56A3.4) clone from the *C. elegans* RNAi feeding library constructed by the lab of Dr. Julie Ahringer, available from Source BioScience. To achieve full knockdown of endogenous and partial depletion of transgenic SPD-5, both clones were grown simultaneously and mixed (30% Ahringer, 70% endogenous) prior to plating. Due to the increased knockdown efficiency of the Ahringer feeding clone, incubation times had to be shortend to 12 hs. Quantification of SPD-5 protein knock down was quantified from western blots using the Gel Analyzer function in Fiji [28].

#### Antibodies and Stainings

Stainings were done following standard procedure described before [6]. The polyclonal mouse αPLK-1 antibody was generated in house by injecting 1 mg of purified full length PLK-1 into mice, purified from serum, and used in a dilution of 1:300. Endogenous SPD-5 and SPD-2 were detected using the previously described polyclonal rabbit αSPD-2 antibody (anit-spd-2_NT_Acid, dil. 1:4000) as well as αSPD-5 antibody (anti-SPD-5_mid_Acid, dil: 1:7200). Commercially available (Life Technologies) Goat anti-Mouse-AlexaFluor594 conjugates, Goat anti-Rabbit-AlexaFluor594 conjugate and Goat αRabbit-AlexaFluor647 conjugate were used for detection with a 1:1000 dilution. GFP signal was detected directly using 488 nm illumination. Images were recorded using an inverted Olympus IX71 microscope, 40X NA 1.00 Plan Apochromat oil objective, CoolSNAP HQ camera (Photometrics), a DeltaVision control unit (AppliedPrecision) and the recording software SoftWoRx 5.5. Centrosome intensity analysis was carried out using Fiji. Briefly, centrosomes were detected automatically using the autothreshold function and analyzed subsequently for mean intensity using the analyze particle function. For quantification of PCM localized SPD-2 and PLK-1 staining signal the high centriolar signal was excluded for the quantification, see Figure S2B for detailed procedure.

#### Centrosome Live Imaging

Centrosomes were detected via imaging GFP::SPD-5 on an inverted Nikon TiE microscope with a Yokogawa spinning-disk confocal head (CSU-X1), a 60x water 1.2 NA Nikon Plan-Apochromat objective with 1.5 X Optovar, and a iXon EM + DU-897 BV back illuminated EMCCD (Andor) 63 x 0.26 μm z-stacks were recorded in 30s intervals with 15% laser transmission. Images were also taken using an inverted Olympus IX81 microscope with a Yokogawa spinning-disk confocal head (CSU-X1), a 60x water 1.2 NA UPlanSApo objective, and a iXon EM + DU-897 BV back illuminated EMCCD (Andor). 46 x 0.4 μm z-stacks were recorded in 30s intervals with 15% laser transmission. For experimental comparisons only recordings from the same microscope were used. Maximum projections were generated from z-stacks and background fluorescence was subtracted from the images. Centrosome area and mean maximum fluorescence intensity were measured throughout development of each embryo by selecting centrosomes using the thresholding and Analyze Particles function in Fiji. Centrosome volume was then calculated using the centrosome radius approximated from centrosome area measurements. Total SPD-5 amounts were approximated from centrosome volumes and densities. We multiplied the volumes with concentration corrected intensity values, assuming that the recorded intensities stemmed from the 30% of the total SPD-5 pool labeled with GFP.

#### *In vitro* SPD-5 scaffold assembly

Proteins were purified and assembled into scaffolds *in vitro* as previously described [12]. For a detailed protocol please refer to [29]. In brief, all reactions were carried out in network buffer (25 mM HEPES, pH 7.4, 135 mM KCl, 125 mM NaCl, 0.2 mM ATP, 10 mM MgCl_2_, 1 mM DTT, 0.02% CHAPS, 0.2% glycerol, 0.025 mg/ml Ovalbumin) plus pre-blocked 0.2 μm red fluorescent polystyrene beads (Invitrogen) to aid in finding the focal plane. All proteins and reagents were stored on ice prior to being mixed, aliquoted into multiple tubes, then incubated at 23°C. 2 μl of a particular reaction was spotted onto a non-frosted cover slide, then covered with 18 x 18 mm pre-cleaned hydrophobic cover slips.

Cover slips were cleaned and made hydrophobic using the following steps. First, cover slips were placed in a Teflon holder and submerged in a 1:20 dilution of Mucasol detergent (Sigma) for 10 min with sonication. Second, the cover slips were transferred to 100% Ethanol and incubated for 10 min with sonication. Third, the cover slips were incubated in a 50 % solution of Rain-X (diluted in ethanol) for >30 min, then washed in ethanol, then twice in water. Finally, the cover slips were dried using N2 gas and stored in a desiccation chamber.

#### PLK-1 inhibition in embryos with permeabilized eggshells

PLK-1 inhibition was performed as previously described [12]. Briefly, L4 worms were seeded onto feeding plates containing bacteria expressing *perm-1* dsRNA and incubated at 20°C for 14–20 hours [22]. Worms containing permeabilized embryos were dissected in an imaging chamber (Microwells) containing osmotic support medium (stock solution: 50 ml ESF 921 Insect Cell Culture Medium (Expression Systems) + 5ml Fetal Bovine Serum + 0.77g Sucrose, filter sterilized; this stock solution was then diluted 60/40 in M9 buffer). To inhibit PLK-1^AS^ prior to mitotic entry, the buffer was exchanged for buffer containing 10 μM 1-NM-PP1 (Cayman chemical company, Cat#13330). At the end of each experiment, FM4–64 (Molecular Probes, # T13320) was added to the well to confirm that the imaged embryo was permeable. Imaging was performed as described above, except that 10s intervals and 8% laser power (4.5 mW) were used.

#### Analysis of diffusion using fluorescence correlation spectroscopy (FCS)

FCS measurements and diffusion analysis were carried out as described previously [17]. Briefly, FCS measurements were made on a LSM 780 microscope equipped with a 40X 1.2 NA water immersion objective and an avalanche photodiode (Zeiss, Jena, Germany) at room temperature using 488 nm excitation. The focal volume was calibrated using Alexa 488 dye (Life Technologies), resulting in the following parameters: beam diameter (ω_xy_) = 0.19 μm, structural parameter (S) = 5 and confocal volume (V) = 0.19 fl. Three measurements with a total of 72 s (24 s each) were taken in each embryo at random positions in the cytoplasm (excluding cell membrane, pronuclei, and centrosomes) after a 1s prebleach. Autocorrelation curves were calculated from the intensity profiles obtained from the measurements and then averaged for each embryo. To obtain diffusion coefficients, each averaged autocorrelation curve was fitted within a time range of 500 ns to 1 s with a single component three-dimensional anomalous diffusion model including a free triplet component to account for fluorophore blinking [30,31]. Statistical analysis on diffusion coefficients was done using Wilcoxon rank sum test. Autocorrelation analysis and data plotting were carried out using a MATLAB script developed in our lab.

## Author Contributions

O.W., J.B.W., and D.Z. designed the experiments and wrote the manuscript. O.W. performed the *in vivo* experiments and quantifications of centrosome size and density as well as protein levels from western blots and stainings. D.Z. developed and tested the model. J.B.W. performed the in vitro experiments and in vivo analysis of centrosome size in the *plk-1^as^* embryos. Y.L.W. and K.O. created the *plk-1^as^ gfp::spd-5* strain. A.S. immunostained embryos. A.A.H and F.J. provided critical feedback for the manuscript.

## Acknowledgements

We thank the Protein Expression & Purification and Light Microscopy facilities of MPI-CBG (Dresden). This project was funded by the Max Planck Society and the European Commission's 7th Framework Programme grant (FP7-HEALTH-2009-241548/MitoSys) and a MaxSynBio grant to A.H. and F. J. J.B.W. was supported by an EMBO fellowship and MaxSynBio.

## Supplementary Information

### Quantitative model of SPD-5 incorporation

We use a theoretical approach to discuss the effects of the nonphosphoraylatable SPD-5^4A^ mutant on the dynamics of PCM assembly and centrosome growth. We extend the previous physical model of centrosome assembly that is based on the physics of liquid droplets [1]. This model can quantitatively account for the volume growth of centrosomes *in vivo*. Here, we extend this model to a situation where in addition to the wild-type form SPD-5^WT^ also the mutant form SPD-5^4A^ is expressed. The biochemical effects of the mutations are not known. By comparing theoretical predictions to experimental data, we aim to identify the biochemical properties of SPD-5^4A^ during centrosome assembly.

Our original model is based on the idea that SPD-5 molecules exist in two different forms [1]. We distinguish the form *A* that is soluble in the cytoplasm and the form *B* that tends to aggregate and phase separate from the cytoplasm. This form *B* is thus the basis of the PCM phase that forms the centrosome. The two forms of SPD-5 can be converted into each other by chemical processes such as phosphorylation. We showed that the conversion of *A* to *B* must be autocatalytic to account for the sigmoidal centrosome growth curves and the reliable initiation of PCM accumulation at the centrioles [1].

We extend our original model by distinguishing wild-type SPD-5^WT^ from mutant SPD-5^4A^. Potentially, both species can exist in the *A* and the *B* form and we thus introduce four molecular species: *A*^WT^, *B*^WT^, *A*^4A^, and *B*^4A^. The dynamics of SPD-5^WT^ and SPD-5^4A^ in the cytoplasm, as revealed by fluorescent correlation spectroscopy (FCS) measurements, are very similar (Figure S4). We thus consider both *A* forms to have the same properties, such that they are both soluble in cytoplasm and can also diffuse in the PCM. The PCM can be formed by aggregation of either *B^WT^* or *B^4A^* or by their combined aggregation (Figures 1B and C and 2C and 1D). We thus infer that they tend to phase separate together in one phase. The PCM volume *V* is thus given by *V* = *N_B_*/*n_B_*, where 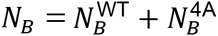 is the total number of molecules in form *B* and *n_B_* is the density of *B* molecules in the PCM. Our data suggests that the density of SPD-5 in the PCM changes only slightly during centrosome growth (Figure 2A). For simplicity, we here consider the case where the density *n_B_* is determined by the physics of phase separation and is constant over time. Below we discuss possible causes for the observed small density variations.

The PCM size increases if molecules of form *B* are generated from their respective *A* forms. Since PCM growth is reaction-limited [1], the *A* molecules can distribute in the PCM more quickly than they get converted to the *B* form. Consequently, the densities 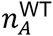 and 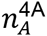 of the *A* forms are approximately homogeneous throughout the PCM. We consider the simple case where the two *A* forms do not interact with each other, such that we can discuss their densities separately. In the simplest case, the equilibrium of the *A* form between the inside and the outside is given by an equilibrium constant *ξ^i^*, which implies that the density 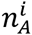 inside the PCM is proportional to the respective density 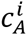 of the *A* form in the cytoplasm, 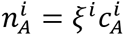 for *i* = WT, 4A. Here, *ξ^i^* can be interpreted as an affinity of the PCM toward molecules of species *A*^i^.

Chemical reactions lead to the creation and removal of molecules of form *B* inside the PCM, which is described by [1]

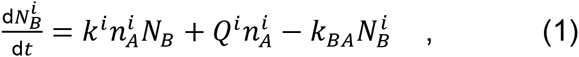

for *i* = WT, 4A. Here, the first two terms describe the production of a single *B* molecule, while the last term accounts for the back-reaction *B* → *A*. The rate-constant *k^i^* in the first term quantifies the rate of the autocatalytic reaction *A + B* → 2*B*. The second term accounts for the catalytic activity of the centrioles, which is quantified by the reaction flux *Q^i^*. Note that this catalytic activity is necessary to initiate centrosome growth [1] and to stabilize two centrosomes in the same cell [2]. We consider the case where both this catalytic and the autocatalytic reaction are driven by the same enzyme, e.g., PLK-1. We thus choose the ratio *N*_0_ = *Q^i^*/*k^i^* of the reaction rates to be the same for the wild-type and the mutant, respectively. Here, *N*_0_ is proportional to the number of catalytic sites on the centrioles.

For the simple case where centrosomes start growing around bare centrioles without PCM, 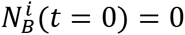, and the concentration 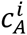 in the cytoplasm remains constant, the number 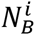 of *B* moleculs in the PCM evolve as

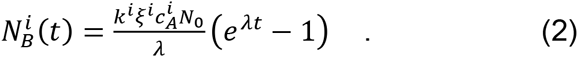

Here,

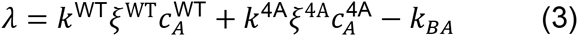

is the PCM growth rate. If we include the depletion of the *A* form in the cytoplasm, 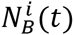 exhibits sigmoidal growth [1]. Since we do not observe a saturation of 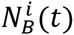 in our experimental data, we focus on the early growth phase where depletion can be neglected.

We compared the predictions of our model to our experimental measurements. In our experiments, we label one of the SPD-5 species with GFP, such that the measured total centrosome fluorescent intensity is a proxy for the total number 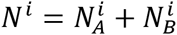 of molecules of species *i*, where 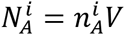, and *V* is the PCM volume. For the case 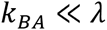, the time evolution of *V* can be approximated by

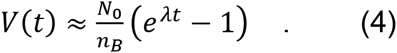

Consequently, the total number of molecules is given by

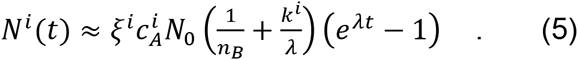

We use this functional form to measure the growth rate *λ* by fitting the function *f*(*t*) = *C*(*e^λt^* − 1) to our measured intensities multiplied by the centrosome volume as a function of time (Figure 4F). Here, the prefactor *C* sets the intensity scale. The measured growth rates were *λ*^WT^ = 0.48 ± 0.08 min^−1^ and *λ*^4A^ = 0.01 ± 0.05 min^−1^ in the experiments with only SPD-5^WT^ and only SPD-5^4A^, respectively. Since 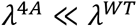, the mutant does not lead to significant PCM growth.

Why does the mutant SPD-5^4A^ not contribute to PCM growth? According to Equation (3) there are three parameters that could be responsible for this: the cytoplasmic concentration could be reduced 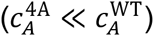, binding to preexisting PCM could be less efficient 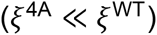, or the transition to the *B* form could be hindered 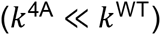.

We measured the total amount of SPD-5 in all experiments (Figure 2G-H and S1). The associated concentrations approximate the cytoplasmic ones in the early growth phase where centrosomes are small. Even though the concentration in the experiment with only SPD-5^4A^ is only half of that of the wild-type, this cannot explain that the growth rates differ by more than an order of magnitude. Consequently, differences in the cytoplasmic concentration cannot explain the observed difference in growth.

We will next show using our fluorescent intensity measurements that the binding coefficients *ξ*^*Ak*^ and *ξ*^WT^ are also not significantly different. Here, we consider the experiments where GFP::SPD-5 in either the wild-type or the mutant form is expressed in a background of unlabeled wild-type SPD-5 (Figure 2C–D). Our measurements (Figure S1A) show that the relative GFP fluorescent intensities in the PCM are similar, which implies similar densities 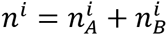 of fluorescent SPD-5 in the PCM (*n*^4A^ ≈ *n*^WT^). Since the mutant does not contribute to PCM growth, the density of its *B* form is negligible 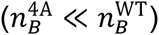. This implies that the density of the mutant *A* form is larger or equal to that of the wild-type 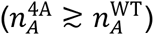. Since the cytoplasmic concentrations 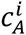 of the fluorescent SPD-5’s are similar (Figure S1) we conclude that the binding coefficients 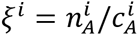 are also similar. In fact, *ξ*^4A^ might even be larger than *ξ*^WT^. Consequently, a reduced binding efficiency of the mutant is not consistent with our data.

The only possible explanation for the mutant’s reduced contribution to PCM growth is that the conversion rate of the mutant from form *A* to form *B* must be much smaller than that of the wild-type (*k*^4A^ ≪ *k*^WT^). Since the mutant cannot be phosphorylated by PLK-1, this furthermore suggests that phosphorylation by PLK-1 plays an important role in this conversion.

Our model implies that the PCM growth rate is proportional to the concentration of SPD-5^WT^, since *λ*^4A^ « *λ*^WT^. We tested this prediction with two additional experiments. First, we quantified centrosome growth in an experiment where both mutant and wild-type SPD-5 were expressed. Using Equation (3), we express the growth rate *λ*^both^ in terms of the already measured growth rate *λ*^WT^ as 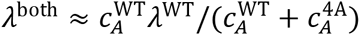. In our experiment, 71% of the SPD-5 molecules are wild-type (Figure S1), which implies *λ*^both^ = 0.71 *λ*^WT^ = 0.34 ± 0.06 min^−1^. This value agrees well with the measured rate 0.31 ± 0.11 min^−1^. Second, we reduced the concentration of SPD-5^WT^ to a fraction of the wild-type concentration without introducing SPD-5^4A^ and found that the PCM volume is reduced by a similar fraction (Figure 2H-J). In summary, our model can capture the growth behavior in all our experiments when the only effect of the mutation of SPD-5 is that it cannot transition from form *A* to form *B*.

In addition to the growth behavior discussed above, we also observed a slight increase in the density of wild-type SPD-5 over time in the experiment without the mutant (Figure 2A). Our current model does not explain this behavior, since it considers phase separation with a constant density *n_B_* of SPD-5 in its *B* form. However, the PCM is not a simple incompressible fluid but a complex polymeric material. Therefore, the incorporation of SPD-5 can lead to a transient buildup of elastic stresses in the PCM, which then relax, e.g., by internal rearrangement of SPD-5 molecules. If this relaxation is slow compared to the SPD-5 incorporation rate *λ*, we expect increased SPD-5 densities over time. Conversely, if the relaxation rate is larger than or comparable to *λ*, the density would be rather constant. We indeed observe these two cases in our experiments (Figure 2A). Our data thus suggests that if only wild-type SPD-5 is present, PCM grows faster than stresses can relax, while they relax during the slower growth in the presence of SPD-5^4A^. Note however that the presence of GFP::SPD-5^4A^ not only affects the growth rate *λ*, but might also have an effect on the relaxation rate, which we cannot measure directly. We therefore did not analyze this further. Nevertheless, these arguments imply a slow stress relaxation rate of less than 0.5 min^−1^, which suggests that the PCM is a rather viscous, gel-like material.

## SUPPLEMENTAL FIGURE LEGENDS

**Fig. S1.**
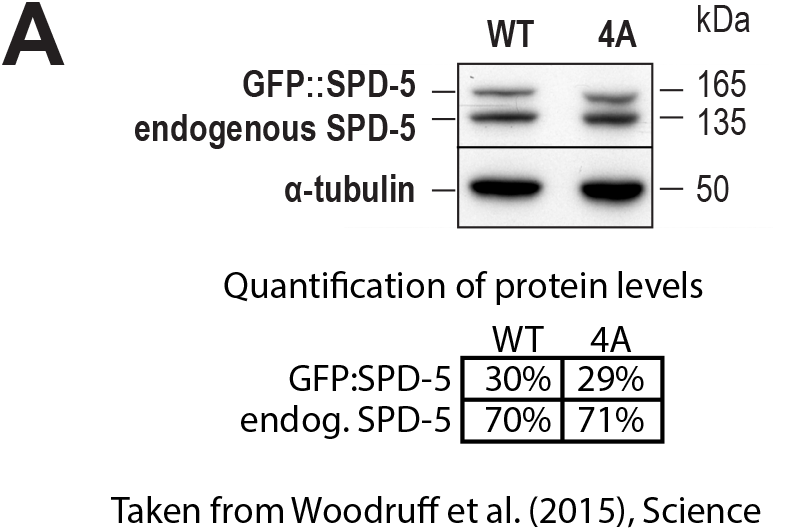
Protein levels in the used transgenic *C. elegans* strains. (A) Western blot analysis of transgenic and endogenous levels of SPD-5 in GFP::SPD-5^WT^ and GFP::SPD-5^4A^ expressing worms as shown previously in [3].

**Fig. S2.**
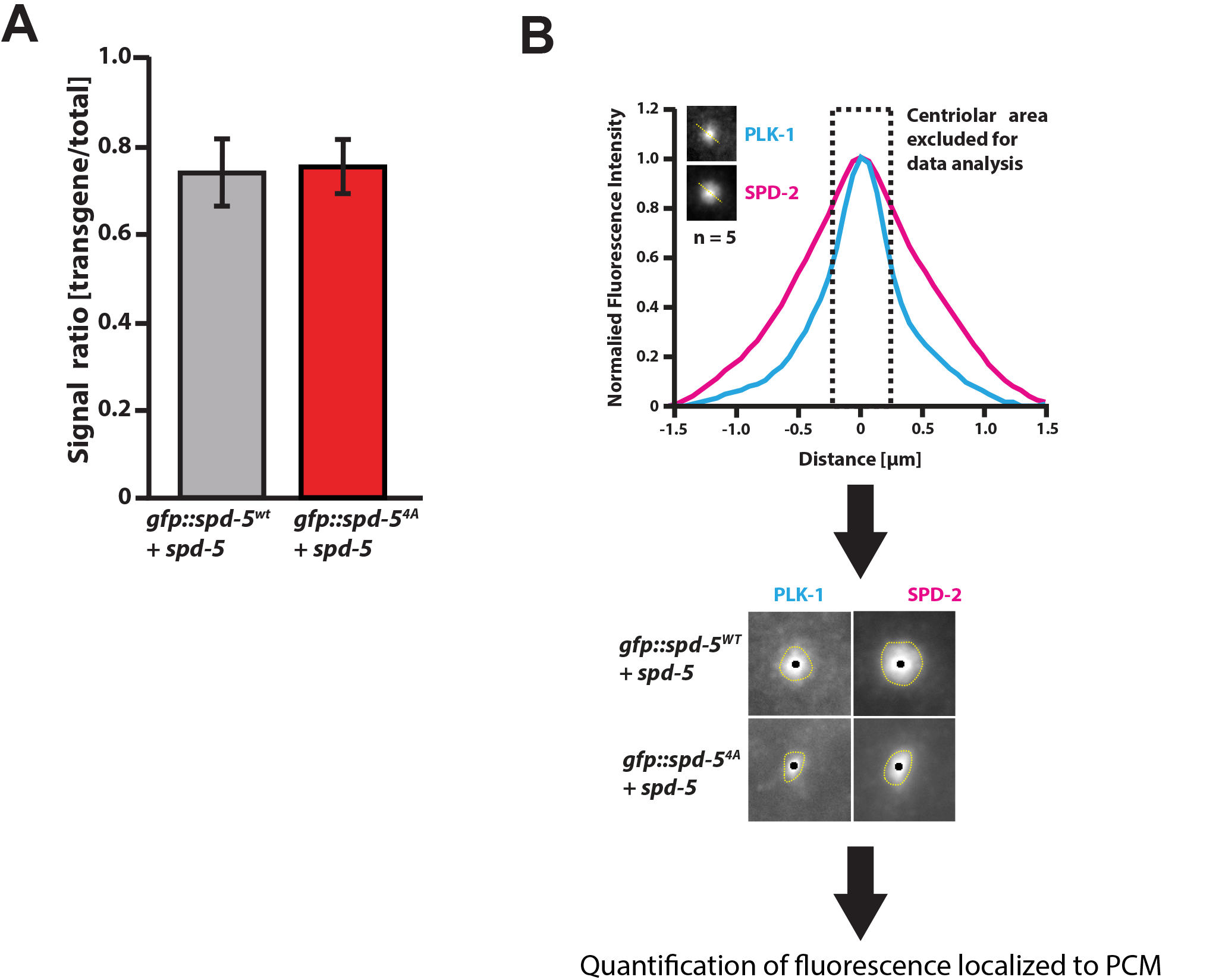
Estimation of protein levels at the centrosome from immunostainings. **(A)** Ratio of transgenic GFP vs. total SPD-5 immunostaining signal at centrosomes based on quantification from Figure 2D. **(B)** Demonstration of the workflow used to quantify PLK-1 and SPD-2 levels at the PCM of GFP::SPD-5^WT^ and GFP::SPD-5^4A^ containing centrosomes (Figure 2F). Top panel shows intensity profiles of PLK-1 and SPD-2 staining signal depicting the strong centriolar localization of the two proteins, which was excluded for the analysis of PLK-1 and SPD-2 levels at the PCM. Bottom panel shows exemplary centrosomes with excluded centriolar signal. Yellow dotted lines indicate the boundary of the centrosome detected through automatic thresholding using Fiji.

**Fig. S3.**
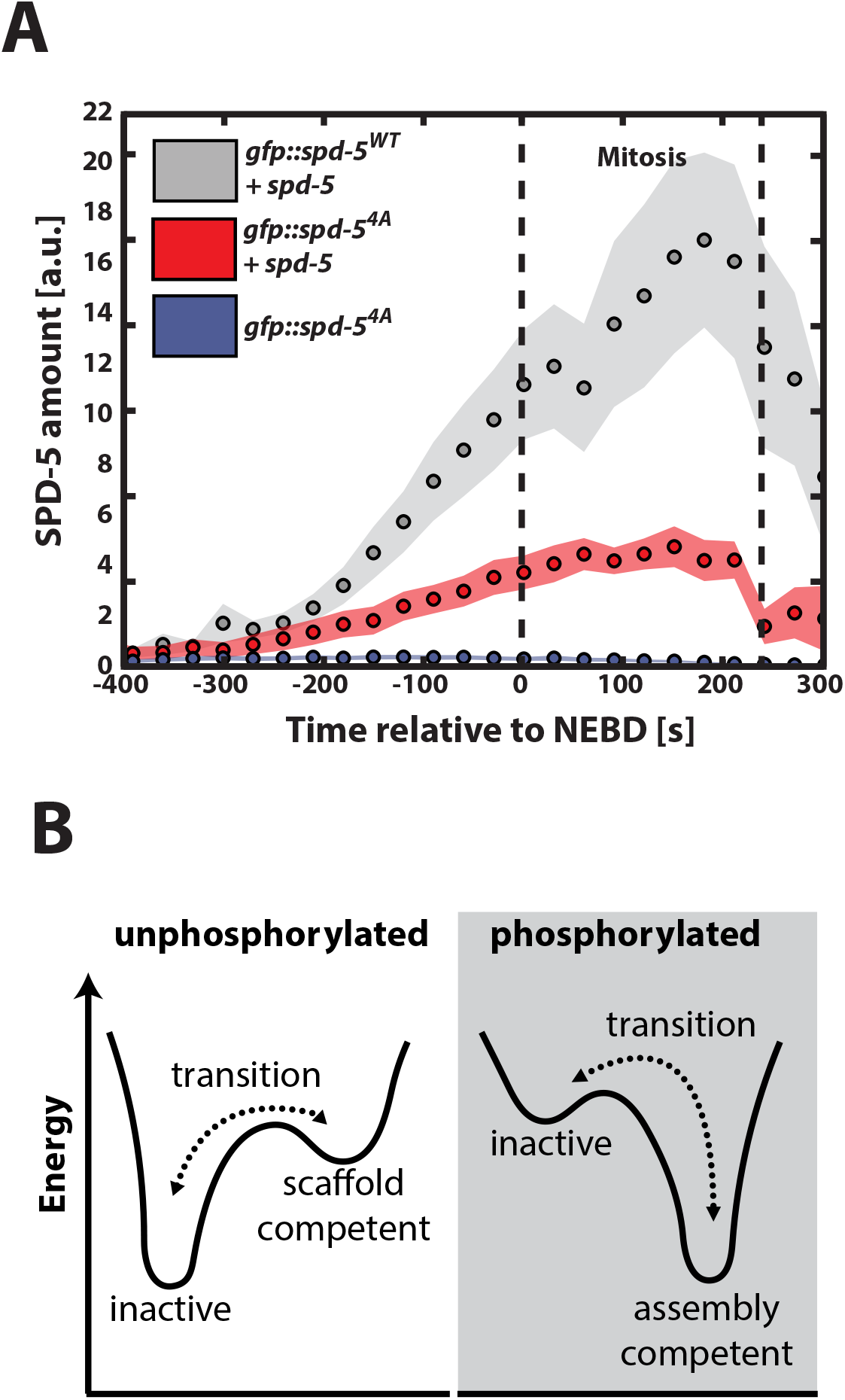
Data and concept used to generate and test the theoretical model. **(A)** Total SPD-5 amount at PCM approximated as the product of centrosome volume (Figure 1D) and mean maximum centrosome intensity (Figure 2A) corrected for protein concentrations. **(B)** Energy diagram of possible conformational states of unphosphorylated and phosphorylated SPD-5.

**Fig. S4.**
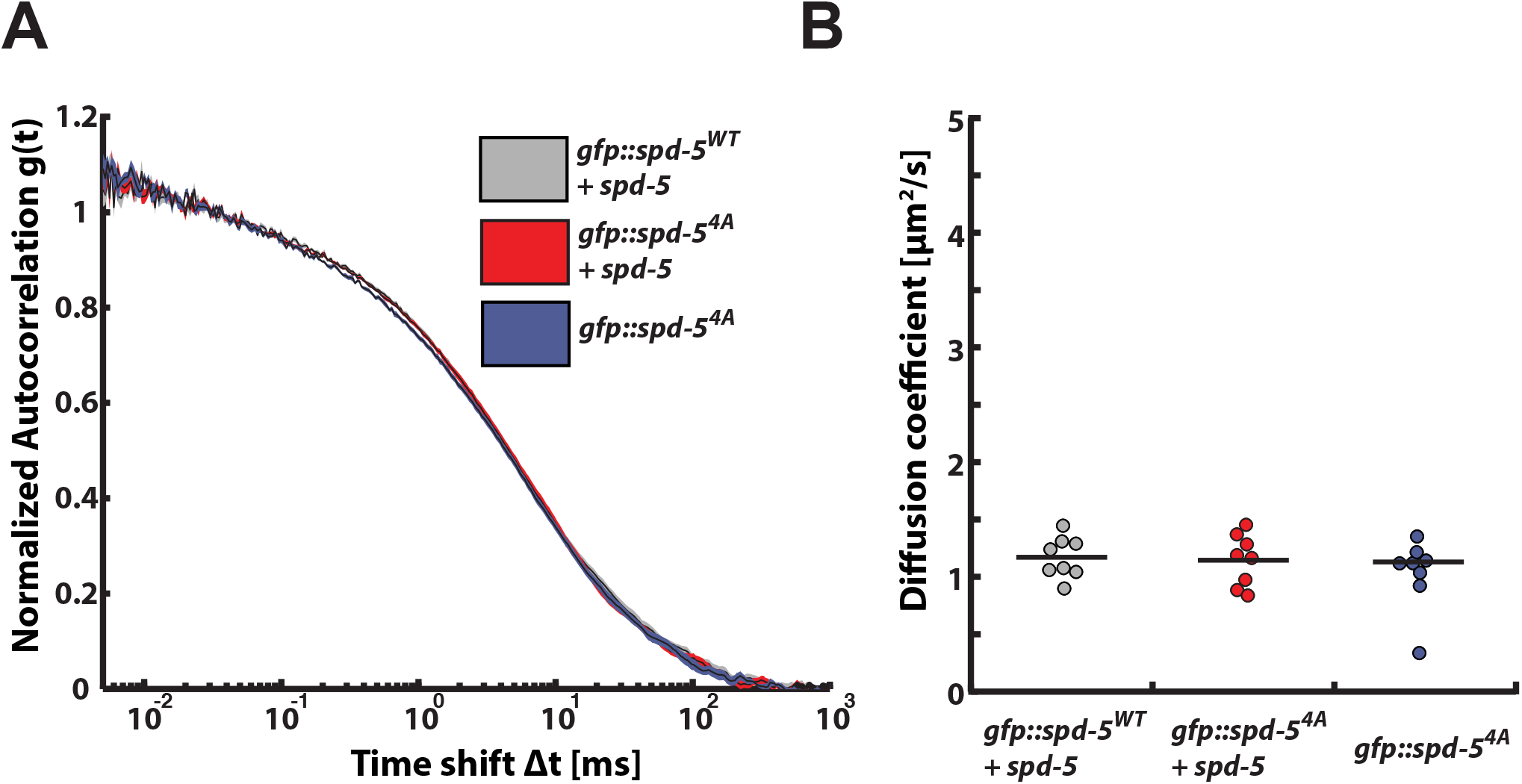
Analysis of cytoplasmic diffusion dynamics of transgenic SPD-5 molecules. **(A)** Normalized autocorrelation curves from fluorescence correlation spectroscopy measurements of GFP::SPD-5^WT^ and GFP::SPD-5^4A^ in the presence or absence of endogenous SPD-5. Lines represent means, shaded areas represent SEM. n = 8 for every experiment. **(B)** Diffusion coefficients obtained from fitting the autocorrelation curves from measurements represented in (A) using a single component 3D anomalous diffusion model as previously described [4]. Diffusion of GFP::SPD-5^WT^ in the presence of endogenous SPD-5 (D = 1.2 ± 0.2 μm^2^/s), and GFP::SPD-5^4A^ in the presence of endogenous SPD-5 (D = 1.1 ± 0.2 μm^2^/s), and GFP::SPD-5^4A^ only (D = 1.0 ± 0.3 μm^2^/s).

